# Reduced beta-hydroxybutyrate disposal after ketogenic diet feeding in mice

**DOI:** 10.1101/2024.05.16.594369

**Authors:** Cody M Cousineau, Detrick Snyder, JeAnna R. Redd, Sophia Turner, Treyton Carr, Dave Bridges

## Abstract

The ketogenic diet (KD) has garnered considerable attention due to its potential benefits in weight loss, health improvement, and performance enhancement. However, the phenotypic responses to KD vary widely between individuals. Skeletal muscle is a major contributor to ketone body (KB) catabolism, however, the regulation of ketolysis is not well understood. In this study, we evaluated how mTORC1 activation and a ketogenic diet modify ketone body disposal in muscle Tsc1 knockout (KO) mice, inbred A/J mice, and Diversity Outbred (DO) mice. Muscle Tsc1 KO mice demonstrated enhanced ketone body clearance. Contrary to expectations, KD feeding in A/J mice did not improve KB disposal, and in most strains disposal was reduced. Transcriptional analysis revealed reduced expression of important ketolytic genes in KD-fed A/J mice, suggesting impaired KB catabolism. Diversity Outbred (DO) mice displayed variable responses to KD, with most mice showing worsened KB disposal. Exploratory analysis on these data suggest potential correlations between KB disposal and cholesterol levels as well as weight gain on a KD. Our findings suggest that ketone body disposal may be regulated by both nutritional and genetic factors and these relationships may help explain interindividual variability in responses to ketogenic diets.

## INTRODUCTION

The ketogenic diet has gained significant popularity in recent years, driven by its potential role in weight loss, improved health outcomes, and enhanced athletic/physical performance. Individuals on a ketogenic diet experience phenotypic variation, encompassing a broad spectrum of responses in weight loss, metabolic adaptation, blood glucose regulation, energy levels, cognitive function, and long-term adherence, which constrains the ability to effectively implement KDs. Ketone body (KB) synthesis, or ketogenesis, primarily occurs in hepatocytes and to a lesser extent in astrocytes or kidney cells [1] The major utilization of ketone bodies happens in the heart, skeletal muscles, and brain [1–3]. The reduction in dietary carbohydrates leads to a decrease in insulin and an increase in glucagon in plasma, promoting hepatic glycogenolysis, gluconeogenesis, and adipose tissue lipolysis.

The regulation of ketogenesis and ketolysis involves a combination of factors including fatty acid flux, hepatic cataplerosis, and the effectiveness of beta-hydroybutyrate (βHB) production and disposal. The rate-limiting enzyme in ketogenesis is 3-hydroxymethylglutaryl-CoA synthase 2 (HMGCS2). It catalyzes the conversion of acetyl-CoA and acetoacetyl-CoA, derived from β-oxidation, into β-hydroxy β-methylglutaryl-CoA (HMG-CoA) and free Coenzyme A (CoA). Mice lacking *Hmgcs2* postnatally exhibited a deficiency in ketogenesis and subsequently developed fatty liver disease spontaneously [4]. However, early weaning proved effective in alleviating hepatosteatosis in postnatal *Hmgcs2* KO mice [4]. In adult mice, insufficient ketogenic activity heightened susceptibility to diet-induced fatty liver disease [4]. Conversely, over expression of Hmgcs2 ameliorated high-fat diet-induced fatty liver disease in adult mice [4]. Together these data show that interruption of ketogenesis affects overall hepatic lipid homeostasis.

In peripheral tissues, KBs, primarily in the form of βHB, re-enter the cell through MCT1-mediated transport. *SLC16A1*, the protein that encodes MCT1, conducts mono-carboxylates, such as lactate, pyruvate, and ketone bodies, across cell membranes [5]. βHBs then undergo re-oxidation to AcAc via BDH1, followed by conversion to Acetyl-CoA by Succinyl-CoA:3-oxoacid CoA transferase (OXCT1 or SCOT) and Acetoacetyl-CoA Thioesterase (ACAT1) for use as ATP a process that contributes to powering muscular work [6–8]. OXCT1 plays a central role in ketolysis in extra-hepatic tissues, activity highest in the heart and kidneys, followed by skeletal muscle and the brain [9]. However, due to skeletal muscle constituting approximately 40% of body mass in adult humans, this organ is the greatest contributor to total KB catabolism at rest [6,10,11]. Although skeletal muscle exhibits a high capacity for KBs, their low circulating concentrations under normal conditions result in a contribution to energy provision in muscle of less than 5% in the post-absorptive state, with free fatty acids being the primary energy source under standard resting conditions [6].

The regulation of ketolysis is not well understood. Prior reports show that PGC1α is necessary and sufficient for promoting ketone body disposal in mice [12]. Exercise also promotes ketone body disposal [13,14], and SLC16A1, BDH1, and OXCT1 are all significantly induced by aerobic exercise training in the MetaMEx database [15]. Among the adaptations to a ketogenic diet in muscle-protein, levels of OXCT1 and ACAT1 were shown to be induced by a ketogenic diet [16], and one report suggests mTORC1 (a positive regulator of PGC1α [17]) is also activated in muscles from mice fed a ketogenic diet [18]. We therefore hypothesized that mTORC1-dependent signaling may cause an adaption in muscles to use ketone bodies in low carbohydrate, high-fat state.

In this study, we evaluated how mTORC1 activation and a ketogenic diet would modify ketone body disposal in mice. We tested this in muscle *Tsc1* KO mice, inbred A/J mice, and Diversity Outbred (DO) mice fed a ketogenic diet, comparing their baseline ketone tolerance tests (KTT) to their post-diet ketone tolerance. We also predicted that variation in ketone disposal may affect cholesterol homeostasis due to the inter-relationship of these metabolic pathways.

## METHODS

### Animal Husbandry and Rodent Diets

Mice were maintained in ventilated cages at 70F at 40-60% humidity with a 12-hour light/dark cycle (ZT0=6:00AM). Mice were provided a normal chow diet (Lab Diet 5L0D; 5% of calories from fat, 24% from protein, 36% carbohydrate) and *ad libitum* access to food and water unless otherwise noted. Muscle-specific *Tsc1* KOs were generated by crossing FVB-Tg(Ckmm-Cre)5Khn/J transgenic mice (RRID: IMSR_JAX:006405) with floxed *Tsc1*^tm1Djk^/J mice (RRID:IMSR_JAX:005680). Mice heterozygous for the floxed allele with or without the cre allele were crossed to generate littermates of KO mice (*Tsc1*^fl/fl^, Ckmm-Cre^Tg/+^) and wild type mice (*Tsc1*^+/+^, Ckmm-Cre^+/+^) as previously described [19]. Wild-type A/J mice (RRID:IMSR_JAX:000646) were purchased at 8 weeks of age from The Jackson Laboratories and at 10 weeks of age were fed control or ketogenic diets for four weeks. Diversity outbred mice (43^rd^ generation, non-siblings RRID: IMSR_JAX:009376) were purchased from the Jackson Laboratories and placed on ketogenic diet at 12 weeks of age for four weeks. For ketogenic diet interventions mice were placed on either a ketogenic diet (Research Diets D17053002, 85% fat, 15% protein, 0% carbohydrates) or a protein-matched synthetic control diet (Research Diets D1053001 10% fat, 15% protein, 75% carbohydrates). Both diets were in meal, not pellet format and were provided in custom jars with holes to provide access. All procedures were approved by the University of Michigan Institutional Animal Use and Care Committee.

### βHB Tolerance Tests

Fed mice were intraperitoneally injected with 1 mg/kg (A/J mouse studies) or 1.5 mg/kg (muscle *Tsc1* KO and diversity outbred studies) of beta-hydroxybutyrate dissolved in PBS at approximately ZT8. Prior to the injection, and then every 15 afterwards, a tail blood draw was collected and analyzed using a Precision Xtra Ketone Body Assay. To summarize these data, we calculated the area under the curve, and baseline-subtracted area under the curve.

### mRNA Analysis

Quantitative real-time PCR was performed by extracting RNA from quadricep muscle lysates using PureLink mRNA kits (Thermo Scientific cat # 12183-018A) and synthesizing cDNA using a high capacity first strand cDNA synthesis kit (Thermo Scientific cat # 4368813). cDNA was amplified using SYBR Green (Thermo-Fisher 4367659) and the primers noted in Table 1 using a QuantStudio 5. Relative expression was determined using the ΔΔCt method. For RNAseq, data was re-analyzed reanalyzed from GSE84312 [19].

### Cholesterol Analyses

Total cholesterol concentration was determined colorimetrically from serum blood samples, using the Infinity Cholesterol Reagent kit (Thermo Scientific cat # TR13421). Following the manufacturer’s instructions, 100 uL of cholesterol reagent and 2uL serum were added to a clear 96-well plate and incubated for 15 minutes at room temperature. Absorbance values were calculated via a standard curve. Each sample was measured in duplicate. The plate was then read on a Molecular Devices Spectramax Plus model plate reader using SoftMaxPro 5.3 at 490 nm.

### Statistical Analyses

All statistical analyses were performed using R version 4.2.2 [20] (RRID:SCR_001905). We set statistical significance for this study at 0.05. For pairwise testing, we first tested for normality via Shapiro Wilk tests, and then equal variance using Levene’s test. On this basis either Mann-Whitney, Welch’s or Student’s t tests were performed as noted in the text and figure legends. For longitudinal data, including βHB tolerance tests we constructed mixed linear models using the lme4 package (version 3.1; [21]). The repeated measure (random intercept) was the animal, and Chi-squared tests were performed comparing models with and without diet terms. For experiments with both rodent sexes, we first tested for modification by sex using the interaction terms from a 2x2 ANOVA, and if significant reported this effect while also stratifying by sex and reporting those results. All raw data and code for this study are available at https://github.com/BridgesLab/TissueSpecificTscKnockouts.

## RESULTS

### mTORC1 Activity Increases the Expression of Ketolysis Genes

While evaluating factors that affect skeletal muscle expression, we noted that several ketolytic genes were elevated in skeletal muscle *Tsc1* KO mice. These mice, due to the deletion of the negative regulator TSC1, have elevated mTORC1 signaling [19,22,23]. We evaluated expression of the putative βHB transporter Slc16a1; which was upregulated (4.0 fold, p_adj_=1.56 × 10^-10^). Among the ketone body catabolic enzymes, Bdh1 (2.1 fold, p_adj_=0.011) and Oxct1 (1.5 fold, p_adj_=1.5 × 10^-5^) were elevated. Acat1 was unchanged in these muscles. These data support the hypothesis that there is transcriptional upregulation of ketolytic genes in muscles with elevated mTORC1 activity.

### Activation of mTORC1 promotes Ketone Disposal

To test whether activation of mTORC1 in muscle tissue alters disposal of ketone bodies, we performed a βHB tolerance test in *Tsc1* KO mice. The Ckmm-Cre induced ablation of *Tsc1* causes activation of mTORC1 in muscle tissues. As shown in Figure 3A, both male and female KO mice cleared the injected beta-hydroxybutyrate much more rapidly than their wild-type littermates. Using mixed-linear models using sex as a covariate, and the animal as a random intercept, we found a significant reduction in βHB levels after the challenge (p=0.004). Similarly, when calculating the area under the curve from 0 to 60 minutes, there was a 25% reduction in the KOs, after adjusting for sex (p=0.016). When stratifying by sex, KOs had 41% lower AUC in males and 11% lower in females, though this modification by sex did not reach statistical significance (p=0.20). Because there is a lack of consensus among studies [18,24,25] evaluating the effect of ketogenic diets on muscle mTORC1 function and suggested activation of this signaling pathway, we evaluated how ketogenic diet feeding alters ketone disposal.

### KD Feeding Does not Improve βHB Disposal in A/J Mice

We hypothesized that prolonged exposure to elevated ketone body levels would lead to physiological adaptations, resulting in improved disposal of ketone bodies. To test this hypothesis, we fed 10-week-old male, wild-type A/J mice a control or a ketogenic diet for three weeks and then performed a KTT. As expected, baseline ketone body levels were elevated from 0.43 mg/dL to 0.75 mg/dL in this assay (p=0.017). Upon injection we were surprised to observe that ketone body levels remain elevated more-so in the ketogenic diet-fed mice than the control mice (Figure 2A), even after subtracting for baseline differences (Figure 2B-C). These data suggest that ketone disposal is not improved after three weeks of a ketogenic relative to a control diet and may actually be somewhat worsened in A/J mice (p=0.274 via linear mixed models).

To understand this surprising lack of adaptation, we performed mRNA quantification of the expression of key transporters and enzymes involved in ketolysis in the quadriceps from these A/J mice. As shown in Figure 2D, most ketolytic gene expression was reduced. Among the MCT1 family transporters that conduct ketone bodies into cells, *Slc16a1* transcripts were modestly reduced in muscles from A/J mice fed ketogenic diets (17% reduced, p_diet_ from a 2x2 ANOVA =0.064). Bdh1 encodes beta hydroxybutyrate dehydrogenase, the interconversion step between beta-hydroxybutyrate and acetoacetate. We found that this gene is downregulated in A/J muscles by 42% (p_diet_ from a 2x2 ANOVA =0.009). *Oxct1* encodes succinyl-CoA:3-ketoacid CoA transferase (SCOT) and was down regulated in both male and female mice by 52% (p_diet_ from a 2x2 ANOVA=8.4 × 10^-9^). While there was no significant sex x diet interaction in these analyses, male mice had 15-30% higher expression of each of these genes compared to female mouse muscle lysates (p_sex_ from a 2x2 ANOVA=4.7 × 10^-3^ for *Slc16a1*, 8.9 × 10^-3^ for *Bdh1* and 0.055 for *Oxct1*). This may suggest enhanced ketone body catabolism in male versus female mice, but this was not explored further. The key finding here is that the transcriptional downregulation of genes involved in ketone body disposal in muscle is consistent with reduced disposal of injected βHB. This begs the question, if ketone bodies are at steady state in serum, but disposal is reduced, does that imply that production may also be reduced? To test this, we looked at three ketogenic enzymes in the livers of these mice. We found downregulation of hepatic *Hmgcl* and *Hmgcs2* in the livers of male mice, and *Bdh1* was downregulated in both male and female mouse livers (Supplementary Figure 1, p_sex x diet_ <0.01 for both *Hmgcl* and *Hmgcs1*).

**Figure 1:**
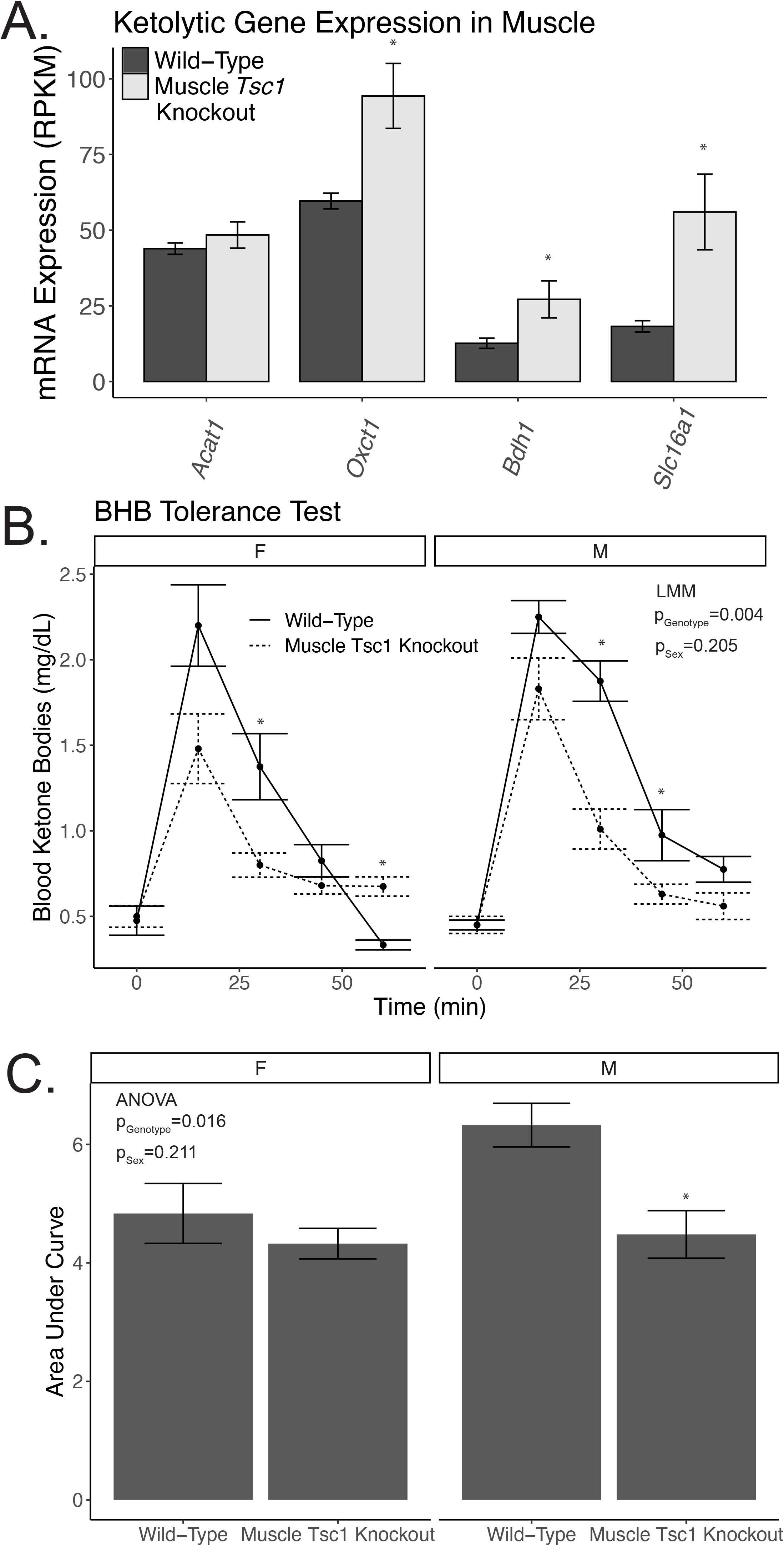
Knockout of Muscle *Tsc1* enhances βHB disposal. A) Ketolytic gene expression in muscle *Tsc1* KO quadriceps. βHB Tolerance tests in male and female wild-type and muscle *Tsc1* KO mice. Mice were injected with 1 g/kg βHB and followed for 1h (n=4-10/group). B) Absolute values stratified by sex. C) Area under the curve. Asterisks indicate p<0.05.

### Evaluation of Diversity Outbred Mice Demonstrates Substantial Variation in βHB Disposal

To evaluate if A/J mice were atypical in their lack of adaptation to improved ketolysis, we performed βHB tolerance tests on diversity outbred mice before or after four weeks of a ketogenic diet. Diversity outbred (DO) mice are genetically unique, so they represent the integrated genetic variability of the intercrossed eight founder strains [26]. As hypothesized by this level of genetic variability, DO mice had variable responses to βHB Tolerance Tests at both baseline and after four weeks of ketogenic diet (Figure 3A). In Figure 3B we describe the within-mouse effects of the diet, again showing substantial between-strain variability, likely due to genetic differences. There was more variability in the area under the curve post-diet than pre-diet (SD_follow-up_/SD_baseline_= 1.93; p<1 × 10^-15^ via Levene’s test), suggesting diet-induced variability. Consistent with our findings using inbred A/J mice, the majority of DO mice had worsened ketone disposal after diet (35 mice), with a smaller number of mice having improved ketone disposal by our assay (10 mice), more than would be expected by chance (p=1.9 × 10^-4^ from a χ^2^ test). Across this cohort there was a significant decrease in ketone disposal (increase in baseline adjusted KTT) in these mice after four weeks of ketogenic diet (p=4.8 × 10^-5^ from a paired Wilcoxon test).

**Figure 2:**
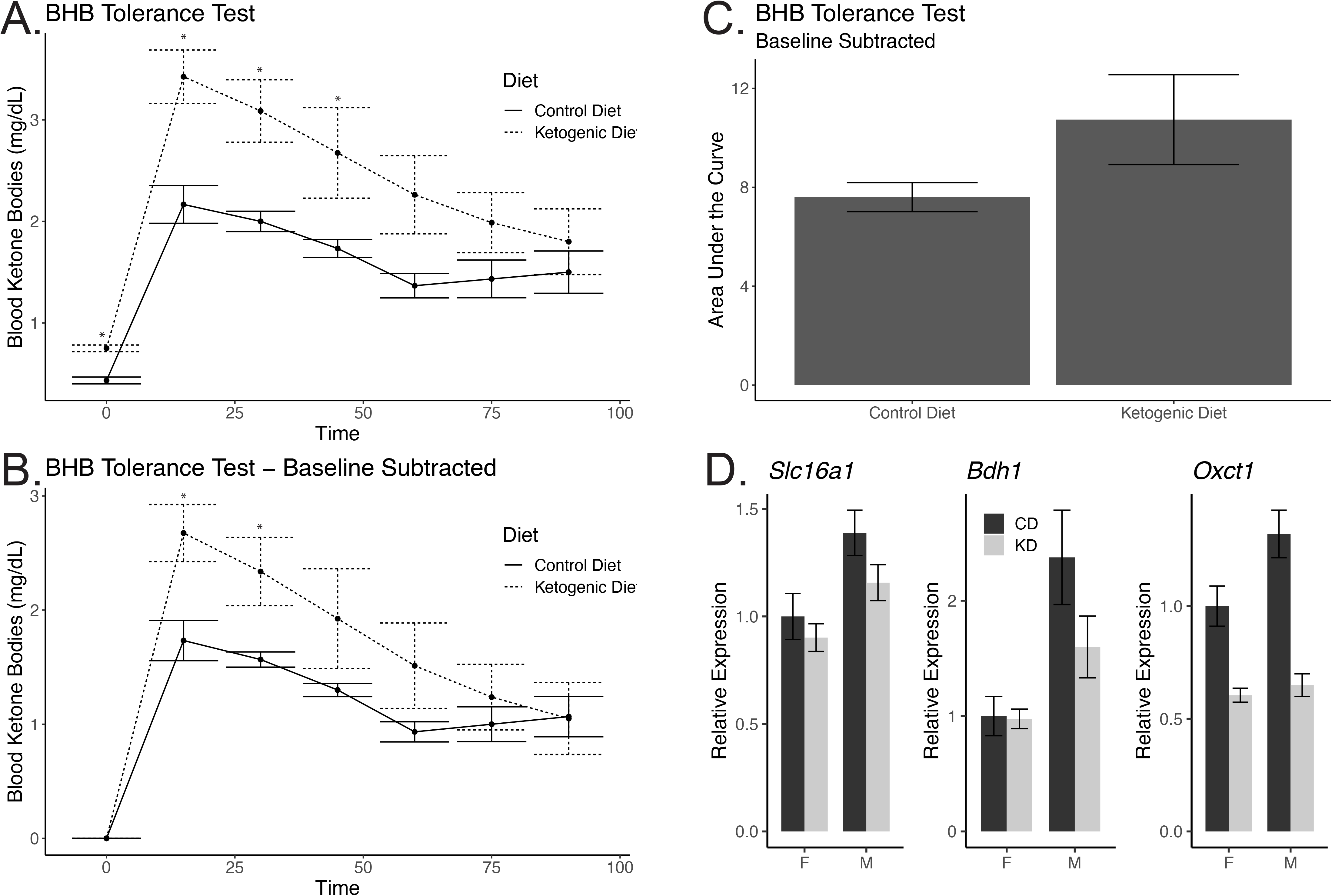
Ketone body disposal is reduced in male A/J mice after ketogenic diet feeding. βHB Tolerance tests in male A/J mice fed a control or ketogenic diet for three weeks. Mice were injected with 1.5 g/kg βHB and followed for 1h. A) Absolute values. B) Baseline subtracted values. C) Area under the curve for baseline subtracted values. D) qRT-PCR analysis of muscle quadriceps mRNA expression in male and female A/J mice. Asterisks indicate p<0.05 from Welch’s t-tests (n=3-8 in A-B and Student’s t-tests in D).

**Figure 3:**
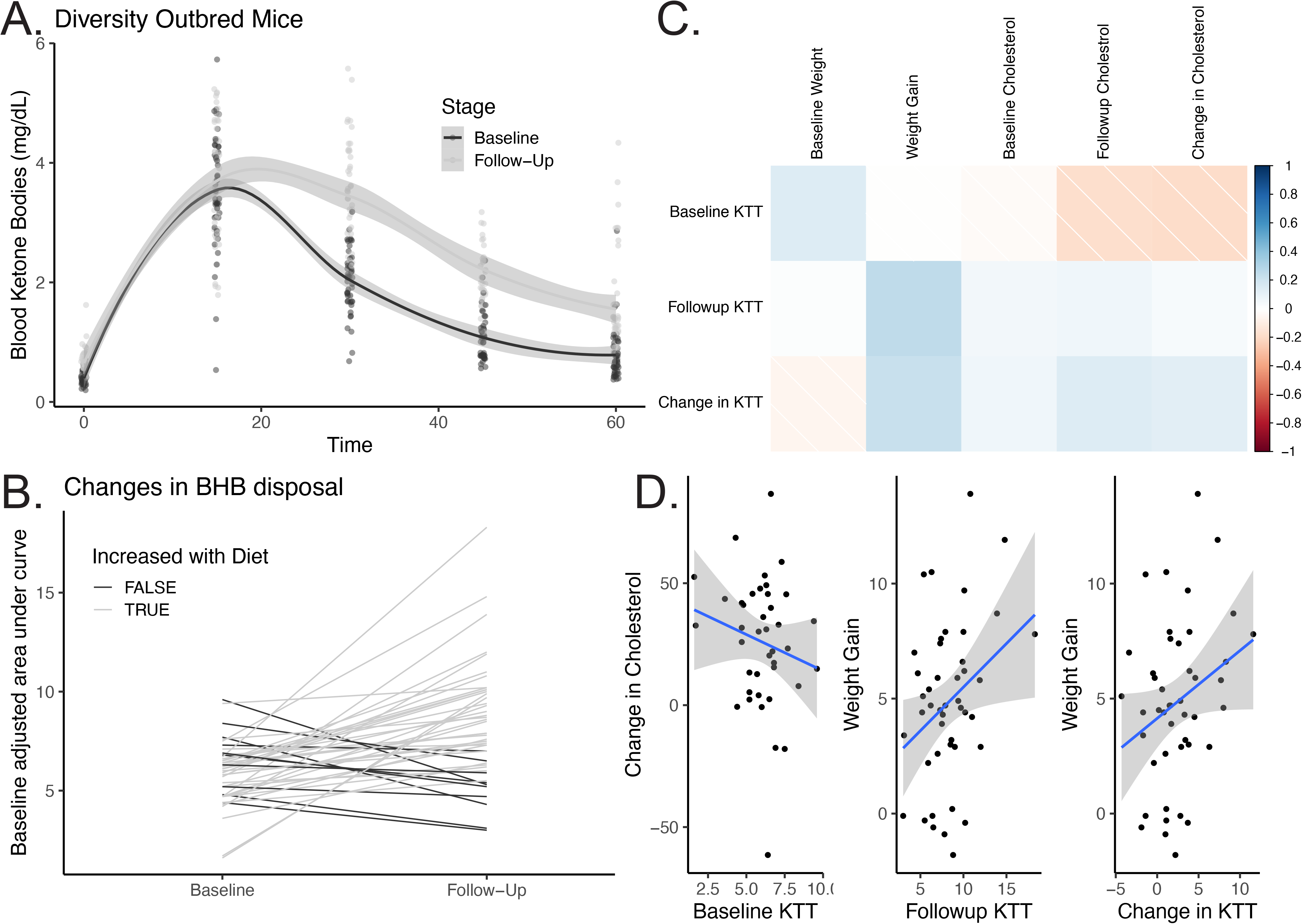
Reduced ketone body disposal in diversity outbred mice after ketogenic diet feeding. A) Ketone tolerance tests at baseline and after three weeks of ketogenic diet in 47 diversity outbred mice B) Individual changes of baseline adjusted ketone tolerance within strains, including whether strains increased or decreased their ketone tolerance. C) Correlation of baseline adjusted ketone tolerance at baseline, follow-up and change over time with weight, weight gain, cholesterol and changes in cholesterol.

To better understand the physiological basis and repercussions of this variation, we performed an exploratory analysis to identify potential correlations between baseline and follow-up changes in ketone disposal, and a variety of other outcomes (Figure 3C). While none of these associations met our threshold for statistical significance, we saw three interesting potential relationships between i) changes in cholesterol and ketone tolerance at baseline (r=-0.18, p=0.26), ii) weight gain on a ketogenic diet and ketone tolerance after the diet (r=0.25, p=0.114), and iii) weight gain and change in ketone tolerance (r=0.23, p=0.158). While this study was underpowered to detect relationships of this size, the findings warrant further investigation into the relationships between ketone disposal and cholesterol and energy homeostasis.

## DISCUSSION

Activation of mTORC1 in skeletal muscle appears to play a positive role in promoting the disposal of ketone bodies and regulating the expression of genes associated with ketolysis. It was tempting to speculate, based on prior reports of mTORC1 activation in muscles after a ketogenic diet, that this was a common adaptation to the low-carbohydrate high-fat (LCHF) state. Surprisingly, our findings revealed that a 4-week feeding period of a ketogenic diet did not result in an enhancement of ketone body disposal in both inbred A/J mice and Diversity Outbred mice, but rather an impairment in disposal of a bolus of βHB. Our results are somewhat at odds with those of Schnyder, et al [16], which showed upregulation of OXCT1 protein in LCHF fed C57BL/6J muscles (BDH1 and SLC16A1 were not tested). In our case we demonstrated downregulation of mRNA, so this could implicate translational regulation, diet, or strain differences with respect to effects on OXCT1, but would not explain our observation of reduced ketone disposal (this was not reported in [16]). In terms of strain differences, we observed that the majority of DO mice experienced a decline in βHB disposal following the dietary intervention (n=35), but there was a subset of mice that exhibited an improvement in βHB disposal as assessed by KTT (n=10).

Given the biochemical relationship between ketone body production, hepatic fatty acid oxidation, and cholesterol biogenesis, it was not surprising to find a potential relationship between these adaptations and cholesterol levels in these mice. The reduction of ketone body disposal may be due to saturation kinetics in skeletal muscle. Prior research has demonstrated that the KB concentration-oxidation relationship may reach a saturating point between 1 and 2 mmol, as evidenced by studies involving fasting of various durations [6] or step-wise βHB infusion [14].

Prior research has shown that knockout of *Ppargc1a* results in modestly lowered expression of BDH1, OXCT1 and ACAT1 in skeletal muscle, brain, and kidneys, along with decreased expression of MCT1 SLC16A1 [12]. This was concordant with impaired ketone body disposal by a similar assay [12]. The converse was also true with muscle transgenic overexpression of PGC1A resulting in induction of Bdh1, Oxct1, Slc16a1, and Acat1 and improved ketone body disposal [12]. It is possible that PGC1α/β is downstream of mTORC1 in muscle. In the muscle *Tsc1* KO data we describe a 34% reduction in *Ppargc1a* mRNA, but a 1.5x increase in *Ppargc1b* mRNA. Because the downstream PGC1a targets are more relevant, we used Harmonizome to identify canonical Ppargc1a targets from ENCODE ChIPseq datasets and contrasted those with differentially expressed genes from muscle *Tsc1* KO mice. Only 4 of28 of these genes were differentially expressed in muscles, lower than what would be predicted by chance.

As it relates to other aspects of physiology in the carbohydrate reduced state, this was to our knowledge the first report of diversity outbred mice and ketone disposal analysis, and we evaluated if this was associated with other relevant traits. Our results of lowered ketone body disposal suggested a reduction in ketogenesis over this period, and given the biochemical overlap between cholesterol synthesis and ketone body synthesis pathways it is not surprising that impaired ketone disposal might be associated with elevated cholesterol, but more data is needed to be confident of this hypothesis.

There are several limitations to the present study. While this study was done in a diverse set of mice, it is plausible that ketone disposal may differ in humans, as may their relationships with cholesterol and energy metabolism. Our approach was to investigate the short-term disposal of a supraphysiological bolus of βHB which again may have different kinetics than elevated steady state levels of ketone bodies. Based on our pilot study of 43 mice we estimate this study is only powered to be able to detect correlations of r>0.55 between two correlated but independent variables. Finally, we do not yet know the mechanism by which skeletal muscle *Tsc1* ablation or mTORC1 activation causes transcriptional changes in ketone body disposal. In our case our assay measured total body ketone disposal, so we do not know that ketone disposal is primarily or exclusively modified in skeletal muscle or also in other tissues.

These findings underscore the necessity for continued in-depth and comprehensive exploration into the factors influencing KB catabolism, including its potential associations with elevated cholesterol. Ketolysis may be influenced by a range of variables, including genetic predispositions, metabolic states, and environmental factors. Based on this research, further investigations are warranted to dissect the molecular pathways, signaling cascades, and genetic factors that contribute to the observed variations in ketone disposal, with the ultimate goal of refining our understanding of the interplay between mTORC1 activation and ketogenic dietary interventions. Such insights may pave the way for personalized therapeutic strategies that consider individual differences in responding to interventions aimed at modulating ketone metabolism.

## ACKNOWLEGEMENTS

This work was funded by R01 DK107535 from the NIH (DB), and Pilot and Feasibility Grants from The Jackson Laboratories and the Michigan Diabetes Research Center (RRID:SCR_015112, supported by P30 DK020572, awarded to DB) and core services provided by the Michigan Nutrition and Obesity Research Center (RRID:SCR_015457, supported by P30 DK89503) and Mouse Metabolic Phenotyping Center in Live Animals (supported by U2C DK135066). TC was supported by the University of Michigan Undergraduate Research Program (UROP). We would also like to thank Dr. Charles Burant and members of the Bridges Laboratory for helpful discussions over the course of this project.

## Supplementary Figures

**Supplementary Figure 1:** Downregulation of liver ketogenic gene mRNA levels after four weeks of ketogenic diet in A/J mice. Asterisks indicate pairwise p<0.05 by Student’s t test (n=7-8/group).

