## Supplementary figures and images for "Reduced beta-hydroxybutyrate disposal after ketogenic diet feeding in mice"

### Supplementary Figure

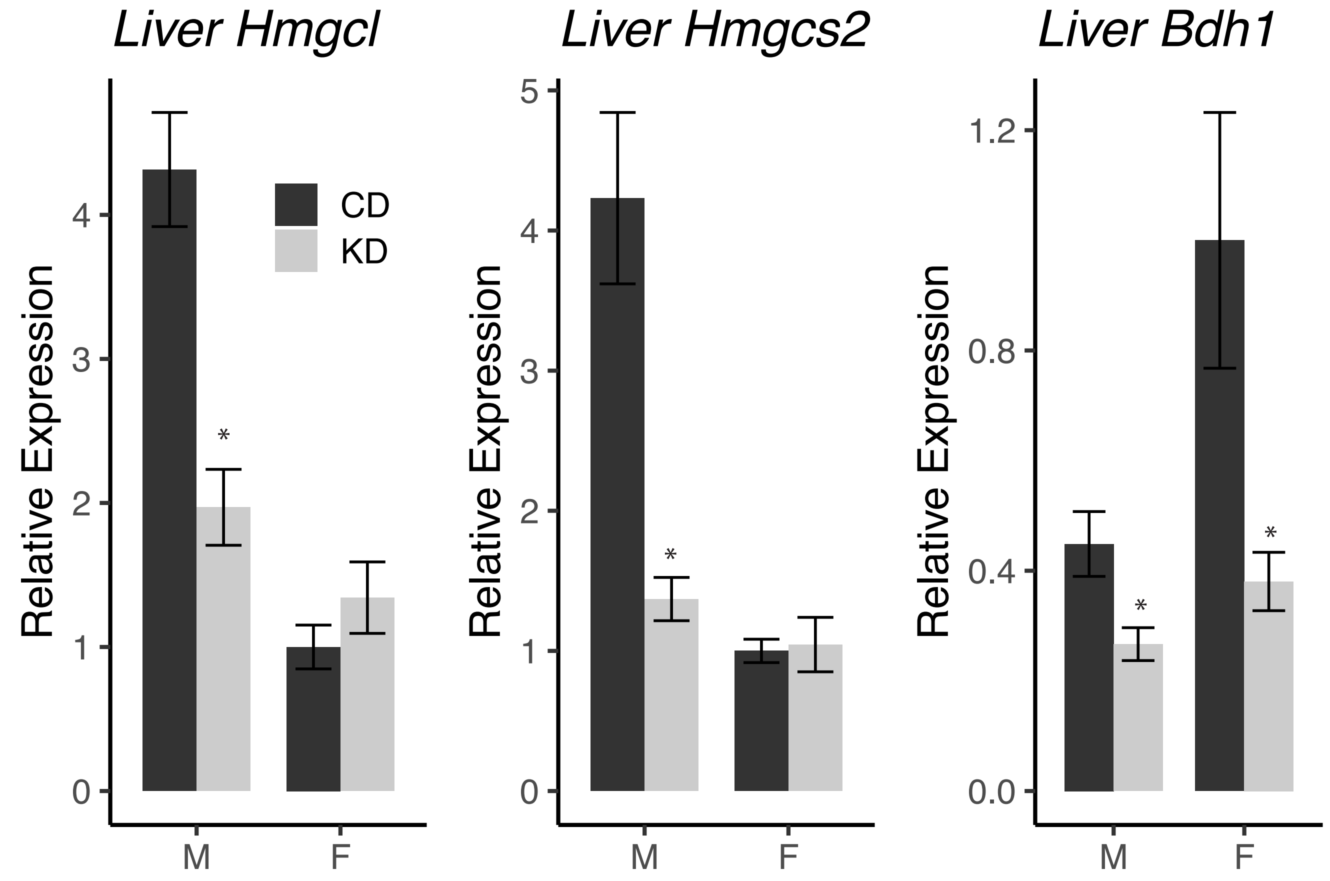
